# Widespread alteration of protein autoinhibition in human cancers

**DOI:** 10.1101/450296

**Authors:** Ashwani Jha, Jennifer M. Bui, Dokyun Na, Jörg Gsponer

**Affiliations:** Michael Smith Laboratories, University of British Columbia.; Department of Biochemistry and Molecular Biology, University of British Columbia.; School of Integrative Engineering, Chung-Ang University, 84 Heukseok-ro, Dongjak-gu, Seoul 156-756, Republic of Korea

**Keywords:** Allostery, autoinhibition, protein regulation, cancer fusions, cancer missense mutations

## Abstract

Autoinhibition is a prevalent allosteric regulatory mechanism in signaling proteins as it prevents spurious pathway activation and primes for signal propagation only under appropriate inputs. Altered functioning of inhibitory allosteric switches underlies the tumorigenic potential of numerous cancer drivers. However, whether protein autoinhibition is altered generically in cancer cells remains elusive. Here, we reveal that cancer-associated missense mutations and fusion breakpoints are found with significant enrichment within inhibitory allosteric switches across all cancer types, which in the case of the fusion breakpoints is specific to cancer and not present in other diseases. Recurrently disrupted or mutated allosteric switches identify established and new cancer drivers. Cancer-specific mutations in allosteric switches are associated with distinct changes in signaling, and suggest molecular mechanisms for altered protein regulation, which in the case of ASK1, DAPK2 and EIF4G1 were supported by biophysical simulations. Our results demonstrate that autoinhibition-modulating genetic alterations are positively selected for by cancer cells, and that their study provides valuable insights into molecular mechanisms of cancer misregulation.

## MAIN

Significant efforts in the cancer research community are underway in order to map all somatic molecular alterations in tumor cells such as single nucleotide variants (SNVs), copy number variations or gene fusions that initiate and drive cancer development^1–6^. The targeted map covers a large number of cancer types and tissues including brain, lung, liver, pancreas, ovary, colon and blood. The currently available map has already revealed a picture of a very complex landscape of genetic alterations^7,8^. Most alterations are likely to be functionally neutral and the result of an increased mutation rate in cancer cells, while a minority of driver alterations provide a selective advantage to cancer cells by helping establish and maintain one or multiple of the six hallmarks of cancer^9,10^. Driver gene fusions, for instance, are responsible for the development of over 50% of all leukemias^11–13^. For the specific case of chronic myeloid leukemia (CML), nearly all of its cases are caused by the chromosomal translocations that give rise to the BCR-ABL1 fusion protein^14^. Establishing functional association between driver alterations and the development of specific cancer types, as it is the case for the BCR-ABL1 gene fusion and CML, has improved biomarker development, diagnostics and clinical care significantly^15–17^.

Equally important as the discovery of drivers is the elucidation of the molecular mechanisms by which the drivers contribute to cancerous cell transformation and progression^8^. For the BCR-ABL1 fusion protein, the transforming potential is, at least in part, due to the disruption of ABL1’s autoinhibition in the gene fusion product^14,18,19^. In autoinhibition, in the most general sense, transient intramolecular interactions between distant parts of the same protein chain reduce or shut down protein function^20–22^. Autoinhibition is a wide-spread form of allosteric protein function regulation, but it is most often seen in signaling proteins. In these proteins, the allosteric regulatory mechanism prevents spurious pathway activation and primes pathways to respond only to appropriate signals. Normally, biomolecular partners and post-translational modifications (PTMs) initiate and reinforce inhibitory intramolecular interactions, or, alternatively, trigger their disruption during signal transmission. As multiple allosteric effectors acting on the same autoinhibited protein can have synergistic or antagonistic effects, autoinhibition has emerged as ideal nexus for complex signal integration and activity-based signal propagation. In cancer cells, this type of allosteric protein activity regulation can be altered due to mutations that remove part of the autoinhibitory switch (e.g. BCR-ABL1 fusion protein) or modulate (strengthen or weaken) the inhibitory intramolecular interactions (e.g. SNVs in the autoinhibited kinase Akt1) ^14,20,23,24^. There has been significant progress and success in the identification of individual proteins that drive cancerous cell transformation due to altered autoinhibition, not least because re-establishing and stabilizing autoinhibition via small compounds has proven to be a successful strategy in cancer drug development^18,25–27^. However, a systematic and robust analysis is needed to reveal whether mutations of allosteric inhibitory switches in cancer is a rare event, or whether altered allostery is highly prevalent and associated with signature molecular changes.

To close this knowledge gap, we integrated cancer genomics data with protein regulation information and systematically analyzed the association between cancer fusions /somatic cancer missense mutations and protein autoinhibtion across multiple tumor types. Our data integration approach reveals a significant enrichment for cancer-associated missense mutations and fusion breakpoints within allosteric inhibitory switches, which in the case of the breakpoints is specific to cancer fusions and not seen in fusions associated with other diseases. Selection for allosteric switches that are recurrently altered by gene fusions or missense mutations across and within cancer types identifies numerous well-established cancer drivers as well as putative new ones. Cancer missense mutations in allosteric switches are associated with specific transcriptional signatures of cancer, which provides insights into the molecular pathways that are activated as a result of altered allosteric protein regulation in cancers.

## RESULTS

### Breakpoints of cancer fusions are enriched within inhibitory allosteric regulatory switches

We sought to exploit recently established maps of genetic alterations, including gene fusions and SNVs, covering various human cancers coupled with data on allosteric protein regulation to examine the association between autoinhibition and cancer (**Fig. 1**). From 12’957 fusion genes collected from the literature (see methods), we identified 5’503 in-frame fusion proteins that were found in cancer cells and had the fusion breakpoint within protein coding regions. We matched these cancer fusion proteins with a set of 234 human proteins for which we found evidence in the literature that their activity is regulation via inhibitory allosteric switches (ASs). The analysis revealed that proteins found in cancer fusions have an enrichment for regulation via autoinhibition when compared to the entire human proteome (Fisher’s exact test, P < 4.42 × 10^−10^)(**Fig. 2A**), proteins that, so far, have not been found in any gene fusion (Fisher’s exact test, P < 1.11 × 10^−16^, and proteins that are part of gene fusions associated with neurodevelopmental disorders or other disease states (Fisher’s exact test, P <0.02) but not cancer (**Supplementary Fig. 1**). As some of these 234 proteins with inhibitory allosteric switches may have been characterized biochemically because they had previously been found in a cancer fusion (e. g. ABL1) and, as a result, their autoinhibition discovered, there may exist a study bias that, at least partially, explains the observed association. Therefore, we analyzed an additional group of 3’381 human proteins that have predicted allosteric regulatory switches (pASs). The predictions were made with an algorithm that we have previously developed and validated for the identification of intrinsically disordered ASs28 (see methods for details). Consistent with the analysis of known autoinhibited proteins, we find that proteins found in cancer fusions are enriched in pAS when compared to the entire human proteome (Fisher’s exact test, P < 3.33 × 10^−16^(**Fig. 2B**) and proteins that have not been found in any gene fusion (Fisher’s exact test, P < 2.22 × 10 ^−16^. As proteins involved in gene fusion have been shown to be enriched in intrinsic disorder, we repeated the analysis only for proteins that contain large disordered regions (IDRs). This control reveals that cancer fusions are also enriched in pAS when only considering proteins that have IDRs (**Supplementary Fig. 2**).

**Figure 1.**
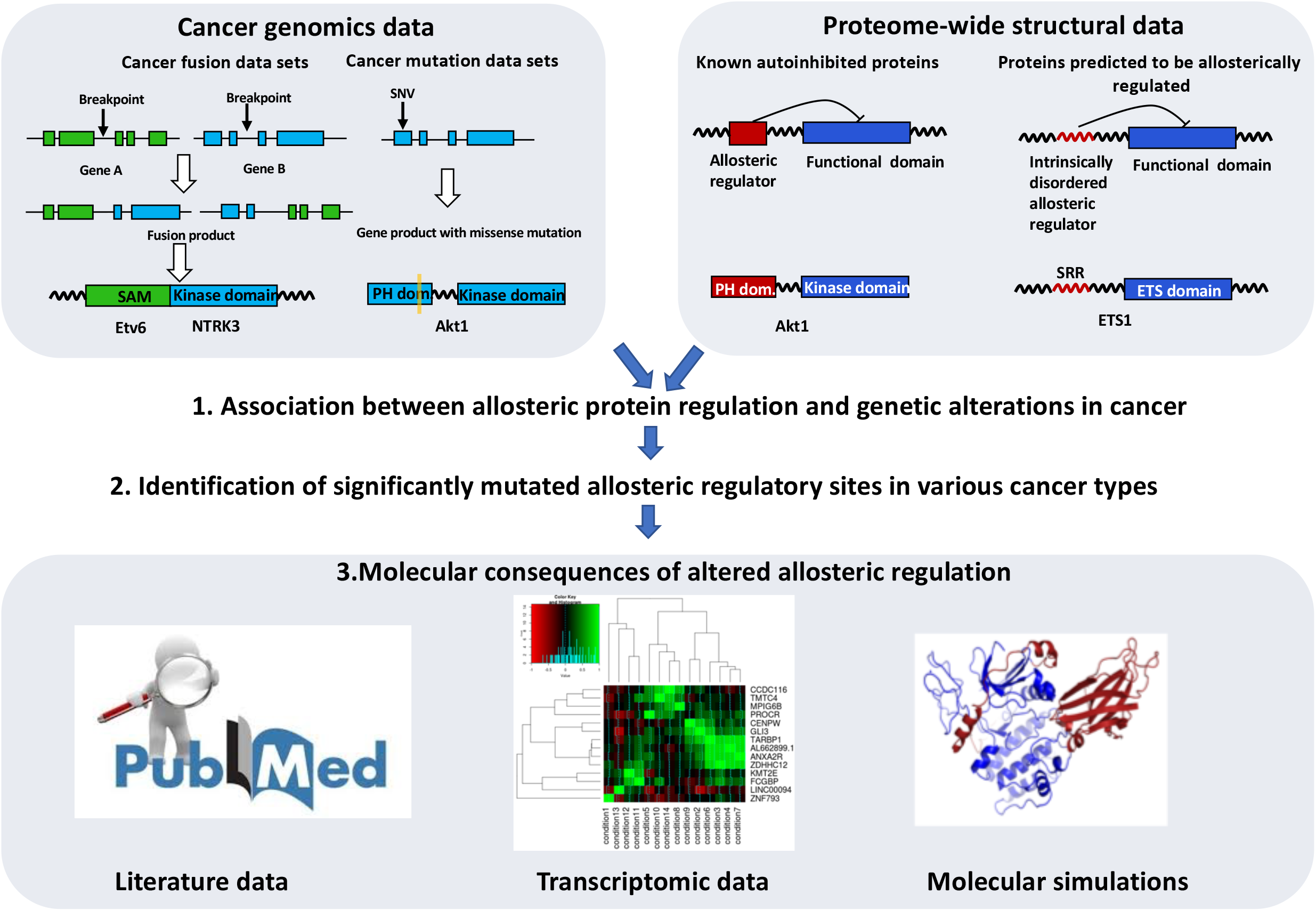
Systematic characterization of altered allosteric protein regulation in cancer. Shown is the flowchart for the integration and analysis of cancer genomics and protein regulation data. In-frame cancer fusion proteins and their breakpoints as well as missense mutations from 33 cancer types are matched with human proteins that have known or predicted allosteric switches (AS). The analysis contains three steps: (1) The enrichment of cancer-associated genetic alterations within AS is assessed. (2) Genes with significantly altered AS are identified. (3) Molecular consequences of altered allosteric regulation are investigated and associated molecular signatures identified.

**Figure 2.**
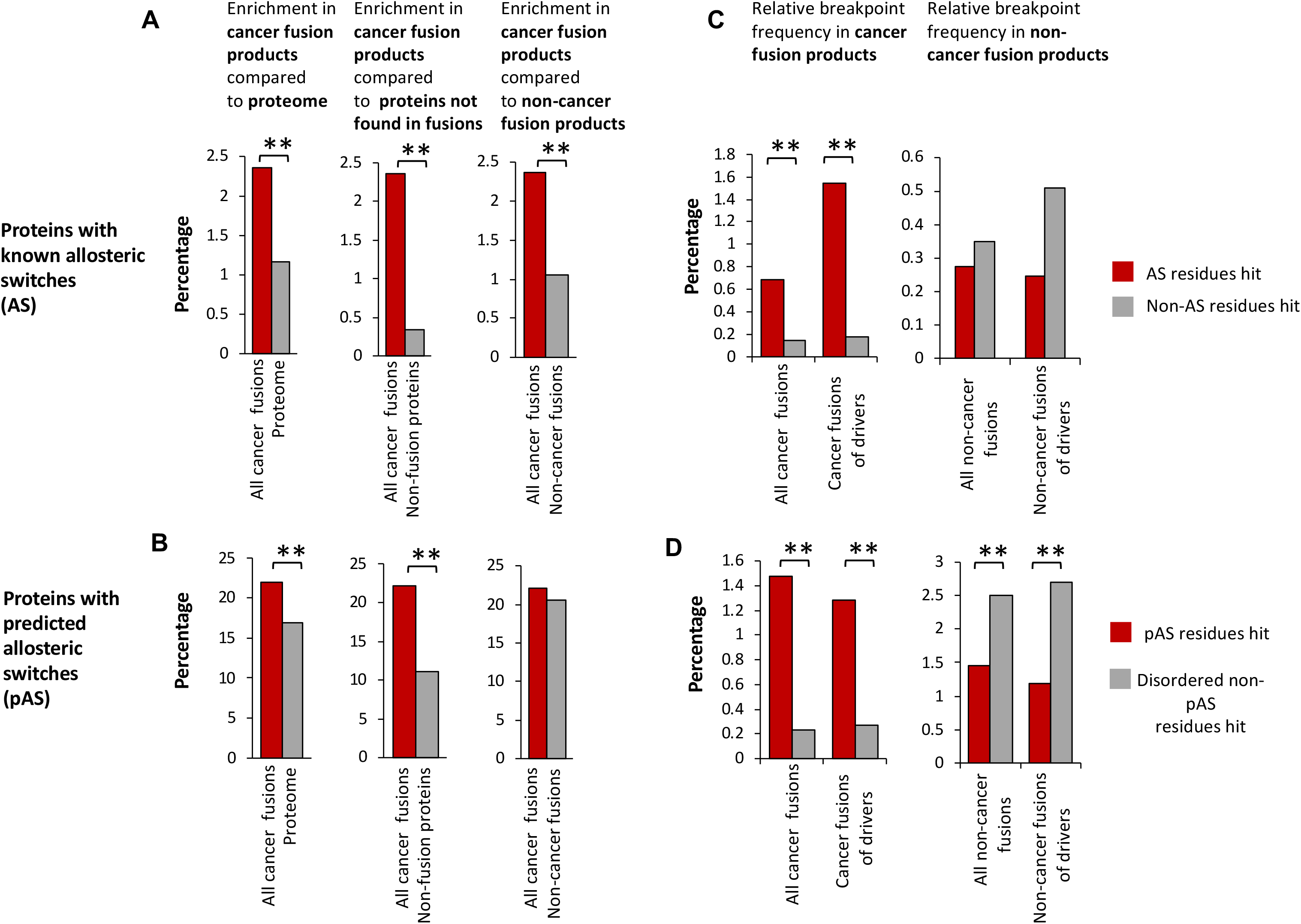
Involvement of allosterically regulated proteins in cancer fusion (A and B) Enrichment of proteins with known (A) or predicted (B) allosteric switches among proteins that are part of cancer fusions (Fisher’s exact test). (C and D) Enrichment of fusion breakpoints within known (C) and predicted (D) allosteric switches. Left, enrichment for all cancer fusions as well as the subset of cancer fusions that involves known driver genes. Right, enrichment for non-cancer fusions as well as the subset of non-cancer fusions that involves cancer driver genes. (Hypergeometric test)

To probe for the relevance of these associations, we analyzed whether autoinhibitory switches get shortened or even removed as result of the gene fusion, i.e., whether breakpoints are located in the AS. Indeed, breakpoints in cancer gene fusions that involve autoinhibited proteins (**Fig. 2C**) are enriched in locations that potentially affect autoinhibition (Hypergeometric test, P < 1.00 × 10^−99^) compared to any other locations in the proteins. This enrichment holds also for the subset of fusions that involve known cancer driver genes (Hypergeometric test, P < 1.00 × 10^−99^). The enrichment appears reversed when considering only gene fusions that are not associated with cancer, even when these non-cancer fusions involve known cancer driver genes (**Fig. 2C**). Dividing driver genes in oncogenes and tumor suppressors reveals that only oncogenes have breakpoints enriched in locations that potentially affect autoinhibition (Hypergeometric test, P < 1.00 × 10^−99^), which is expected as driver alterations in oncogenes are activating while inactivating in tumor suppressors (**Supplementary Fig. 3**). We repeated the breakpoint analysis for proteins with predicted allosteric switches. This analysis reveals that breakpoints are also enriched in pASs when compared to other disordered regions of the same cancer fusion proteins (**Fig. 2D**) (Hypergeometric test, P < 1.00 × 10^−99^) and cancer fusions containing known drivers (Hypergeometric test, P < 1.00 × 10^−99^), respectively. Interestingly, breakpoints are significantly enriched outside of pASs in fusions that are not associated with cancer (Hypergeometric test, P < 3.33 × 10^−64^ and P < 1.10 × 10^−3^, respectively). Consistent with the analysis on known AS, only oncogenes have breakpoints enriched in predicted pASs (Hypergeometric test, P < 2.94 × 10^−34^, **Supplementary Fig. 3)**. We confirmed these enrichment results for pAS in control calculations that defined allostery-altering fusions as those where breakpoints are not only in the pAS but also between the pAS and the protein domain which function is regulated (see methods and **Supplementary Fig. 4)**.

**Figure 3.**
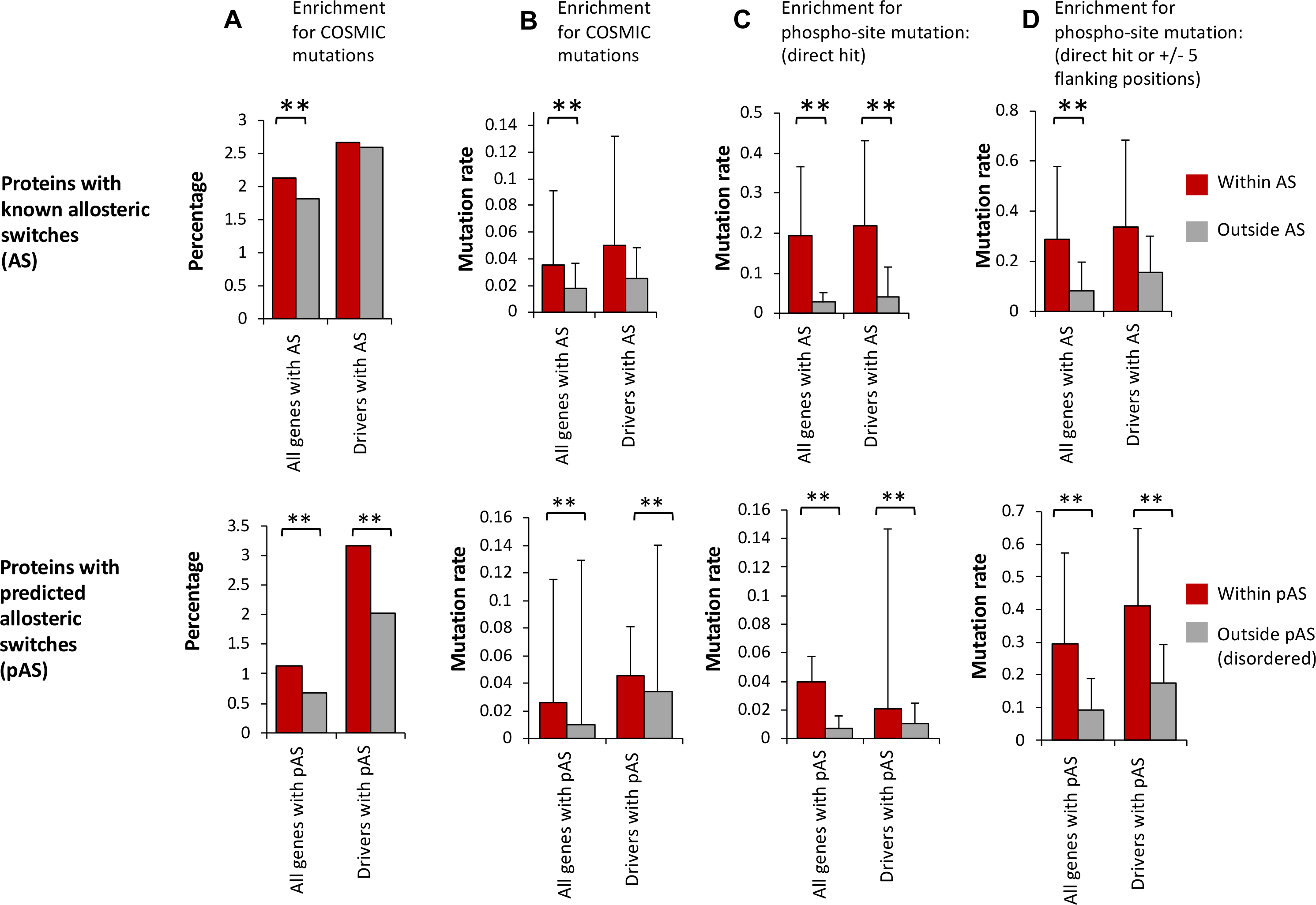
Missense mutations of allosteric switches (A) Percentage of residues within and outside of (p)AS (top AS and bottom pAS) that are hit by missense mutation (hypergeometric test). (B) Averaged mutation rates for residues within and outside of (p)AS. The distributions of averaged mutation rates were compared using Student’s T-test. (C) Averaged mutation rates for phosphorylation sites within and outside of (p)AS. (D) Averaged mutation rates for phosphorylation sites and adjacent residues (±5 residues) within and outside of (p)AS.

### Cancer missense mutations are enriched within inhibitory allosteric regulatory switches

Next, we mapped 7’799 missense mutations from the COSMIC database^29^ on human proteins with AS to determine whether residues in these experimentally-mapped allosteric regulatory switches are frequently mutated in cancers. A significantly higher percentage of sequence positions in ASs have at least one cancer-associated mutation when compared to other locations in the same protein (**Fig. 3A**) (Hypergeometric test, P < 4.10 × 10^−4^). As some sequence positions are mutated multiple times, we also computed the average mutation rate, and found the rate significantly higher inside the ASs compared to outside (**Fig. 3B**) (T-test, P< 0.04). Recent reports have highlighted the potential importance of cancer-related mutations close to or at phosphorylation sites^30,31^. To control for this possible co-variant, we calculated the average mutation rate at (**Fig. 3C**) or close to known phosphorylation sites (**Fig. 3D**) inside and outside of ASs. Phosphorylation sites and residues close-by in ASs have a significantly higher mutation rate than the equivalent sites outside ASs (T-test, P< 0.02 and P< 4.09 × 10^−4^, respectively). When the analysis was restricted to ASs in known cancer drivers (**Fig. 3A-D**), similar trends were observed but, probably due to the lower number of cases, only the mutation rate at phosphorylation sides inside ASs of drivers was found statistically higher than outside (**Fig. 3C**) (T-test, P< 0.03).

We repeated this analysis for the 3’381 human proteins with pASs. Consistent with the analysis of proteins with ASs, we find that a significantly higher percentage of residues in pAS of all human proteins or known drivers have cancer-associated mutation (Hypergeometric test, P < 2.02 × 10^−26^ and P < 9.01 × 10^−11^, respectively) (**Fig. 3A, bottom**) when compared to other disordered parts of the same proteins. In addition, the average mutation rate is higher within pASs, at phosphorylation sites or close to phosphorylation sites within pASs of all human proteins (**Fig. 3B-D, bottom**) (T-test, P< 0.02, P< 0.04, and P<1.74 × 10^−14^) or known drivers (T-test, P< 0.02, P< 0.05, and P<1.46 × 10^−14^), respectively, when compared to disordered regions outside pASs. We found similar trends when taking as a reference the mutation rates of all protein regions outside pASs and not only the disordered parts (**Supplementary Fig. 5).**

### Significantly disrupted inhibitory allosteric switches identify known cancer genes and predict new ones

Motivated by the observed associations, we decided to identify those human genes with recurrent fusion breakpoints in regions that would alter the regulatory effect of known or predicted allosteric switches, (p)ASs, (see Methods) and assess their cancer driver potential. Various methods exist that exploit high cancer mutation rates within genes or mutation clustering to identify drivers^3,4,30,32–37^, but these methods often provide no or little insight into how mutations affect protein function and induce cancer formation. Focusing on allosteric switches provides testable hypotheses on the molecular mechanism by which genetic alterations could induce cancerous cell changes. Thus, we calculated P-values of the binomial probabilities of finding k or more fusion breakpoints within regions that encode the extended (p)AS (see methods for details). Only a pan-cancer enrichment analysis (33 cancer types) was possible because of the low number of reported fusion events. We identified 37 genes with significantly disrupted allosteric switches (**Fig. 4A)**. Most of these genes are expected to be oncogenes because truncation or removal of autoinhibitory switches, as a result of fusion, should lead to activation. Indeed, 13 of the 37 genes (ABL1, RET, BRAF, PDGFRA/B, ALK, FGFR1/2/3, MET, ERG, ETV6 and MYB^8^; **Fig. 4B** and **Supplementary Fig. 6**) are well-known oncogenes, which is a significant enrichment compared to the number of known oncogenes found in all cancer gene fusions (Fisher’s exact test, P< 2.14 × 10^−10^) and cancer gene fusions involving proteins with (p)ASs (Fisher’s exact test, P< 5.44 × 10^−7^). Importantly, disruption of autoinhibition is known to contribute to the oncogenic activity of several of these genes including ABL1^14^, BRAF^38^, PDGFRA^39^ and MYB^40^. Interestingly, one of the 37 genes is the tumor suppressor NF-1. Neurofibromin, the protein product of NF-1, shuttles, like many tumor suppressors^41^, between cytoplasm and nucleus^42^. Nuclear import depends on a NLS^42^ in the C-terminus that also contains a pAS. As appropriate cellular localization of neurofibromin is key to its function^43^, it is conceivable that a removal of the predicted pAS (and the NLS) as a result of fusion will affect its tumor suppressor activity.

**Figure 4.**
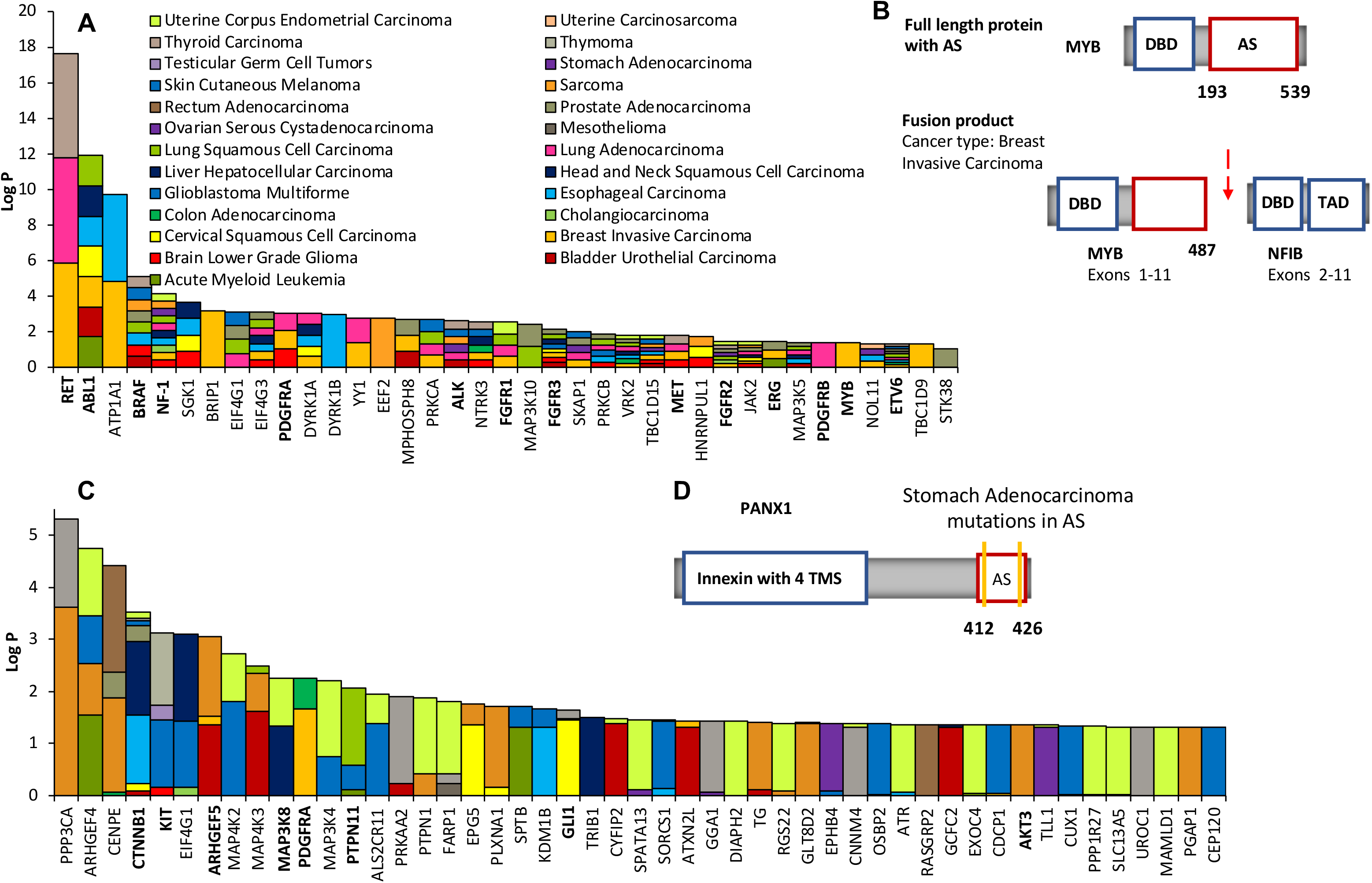
Genes with recurrently altered allosteric switches (A) 39 genes with recurrent fusion breakpoints in their (p)AS across 33 cancer types ranked according to the FDR-corrected P values. Cancers in which breakpoints were found in the (p)AS are indicated by the color. Known cancer driver genes are highlighted in bold. (B) Structural details of the gene MYB and their changes in the MYB-NFIB fusion that is found in breast invasive carcinoma. MYB encodes the transcriptional activator Myb that contains a DNA-binding domain (DBD) and a large C-terminal segment that acts as allosteric switch (AS). In the MYB-NFIB fusion, the fusion breakpoint is located in the AS (highlighted by the red arrow), which, as a result, gets shortened. (C) Top ranked genes with recurrent missense mutations from a specific cancer in their (p)AS. Genes are ranked according to the FDR-corrected P values. Cancer types in which these missense mutations were found are indicated by the color. Known cancer driver genes are highlighted in bold. (D) Structural details of the gene PANX1 with the location of missense mutations in its AS. PANX1 encodes the plasma membrane channel pannexin-1 that contains an innexin domain with four transmembrane segments (TMS) at the N-terminus and an inhibitory allosteric switch at the C-terminus.

We found literature evidence for strong associations with cancer for 19 other genes in the set of 37, highlighting their driver potential. These 19 genes include the known putative drivers SGK1, PRKCA/B, and JAK2, but also new ones such as DYRK1A/B and eIF4G1/3 (**Supplementary Table 1)**. Dual specificity tyrosine-phosphorylation-regulated kinases 1A and B (DYRK1A/B) have been mainly studied in the context of Down syndrome but only recently for their oncogenic potential^44^. Overexpression of DYRK1A as a result of the trisomy 21 has been linked to increased risk of leukemia in children^45^. Moreover, in vitro experiments revealed that the C-terminal tail of DYRK1A contains an autoinhibititory region (**Supplementary Fig. 7A)** and cleavage of this autoinhibitory region by calpain I increase kinase activity in vitro^46^. Thus, removal of the AS, as a result of gene fusion, may constitutively activate DYRK1A and contribute to cancerous cell transformation in certain tissues. Of interest are also the eukaryotic translation initiation factors eIF4G1/3. Overexpression of their interaction partner eIF4E has been found in many solid tumors, where eIF4E promotes the translation of various malignancy-associated mRNAs^47^. The promotion of translation relies on eIF4E’s cap-binding activity and the stimulation of helicase activity of the eIF4A/eIF4G complex. eIF4G1/3 has an AS that inhibits basal helicase activity, and eIF4E is recruited to relieve the inhibitory break and induce full helicase activity (**Supplementary Fig. 7B)**. Moreover, truncation constructs of eIF4G1 that lack the AS have been shown to be able to support efficient translation of capped mRNA^48^. Thus, it is conceivable that removal of the AS in eIF4G1/3 as a result of fusion may promote the translation of malignancy-associated mRNAs^49–52^.

Overall, the 37 genes with significantly disrupted (p)AS across the 33 cancers are significantly enriched in tyrosine kinases (Hypergeometric Test, P< 3.92 × 10^−12^). Nine (RET, PDGFRA, PDGFRB, FGFR1-3, ALK, MET, NTRK3) of them are receptor tyrosine kinases (RTKs), which represents a significant enrichment compared the number of RTKs in all cancer gene fusions (Fisher’s exact test, P< 1 × 10^−99^) and RTKs in cancer fusions involving genes that encode (p)ASs (Fisher’s exact test, P <3.17 × 10^−6^). This enrichment for RTKs is not a surprise because altered autoinhibition in RTKs, as a result of gene fusion, has been found to be a driver in several human cancers. More importantly, it has been demonstrated that shortening of the autoinhibitory juxtamembrane region or the C-terminal tail in RTKs can be sufficient to constitutively activate them and induce cell transformation^39,53^. This said, it is clear that RTKs, like other kinases and transcription factors, often get activated in cancer fusions because of the addition of dimerization domains that promotes activating homodimerization, a mechanism that can even overcome autoinhibition^39,53^. We analyzed how often fusions add a dimerization domain to each of the 37 identified genes. On average, 47% of all fusion events that involve the 37 genes and generate a product with a disrupted/removed (p)AS occur with a fusion partner that encodes a dimerization domain. However, 53% of all fusion events involving these genes result in products that have a disrupted (p)AS and no readily identifiable, additional dimerization domain, which suggests that altered autoinhibition may provide a rational for known or unknown oncogenic activities of these fusion products.

## Significantly mutated inhibitory allosteric switches are associated with molecular signatures of cancer

We also identified human genes with recurrent somatic cancer missense mutations in (p)AS. To this end, we mapped 18’845 somatic missense mutations from NIH-GDC^54^ on genes with regions that encode (p)AS across 33 cancer types and calculated enrichments using a binomial test (see Methods for details). We identified 302 genes with significant mutated (p)AS, some of them in multiple cancer types (**Fig. 4C**). Although genes with recurrently mutated (p)AS are found in 29 different cancer types, several cancers including melanoma and lung adenocarcinoma have a higher burden of genes with recurrently mutated (p)AS (**Fig. 5A**). For those cancer types with more than 10 such genes, an enrichment analysis revealed that these genes have a significant association with cancer pathways (**Supplementary Table 2)**. 29 of the 302 genes with recurrently mutated (p)AS are well-established drivers (**Supplementary Table 3)**, which is a significant enrichment with respect to the number of known drivers in the human proteome (Fisher’s Exact Test, P < 1.45E^−10^) as well as compared to known drivers among proteins with (p)AS (Fisher’s Exact Test, P < 9.58E^−6^). A literature search further provided evidence that 136 of the 302 genes had been associated with cancer in previous studies, which is a significant 21-fold enrichment (Fisher Exact test, P < 2.2 x 10^−16^) as compared to all genes associated with somatic cancer from the COSMIC database (**Supplementary Table 3)**.

**Figure 5.**
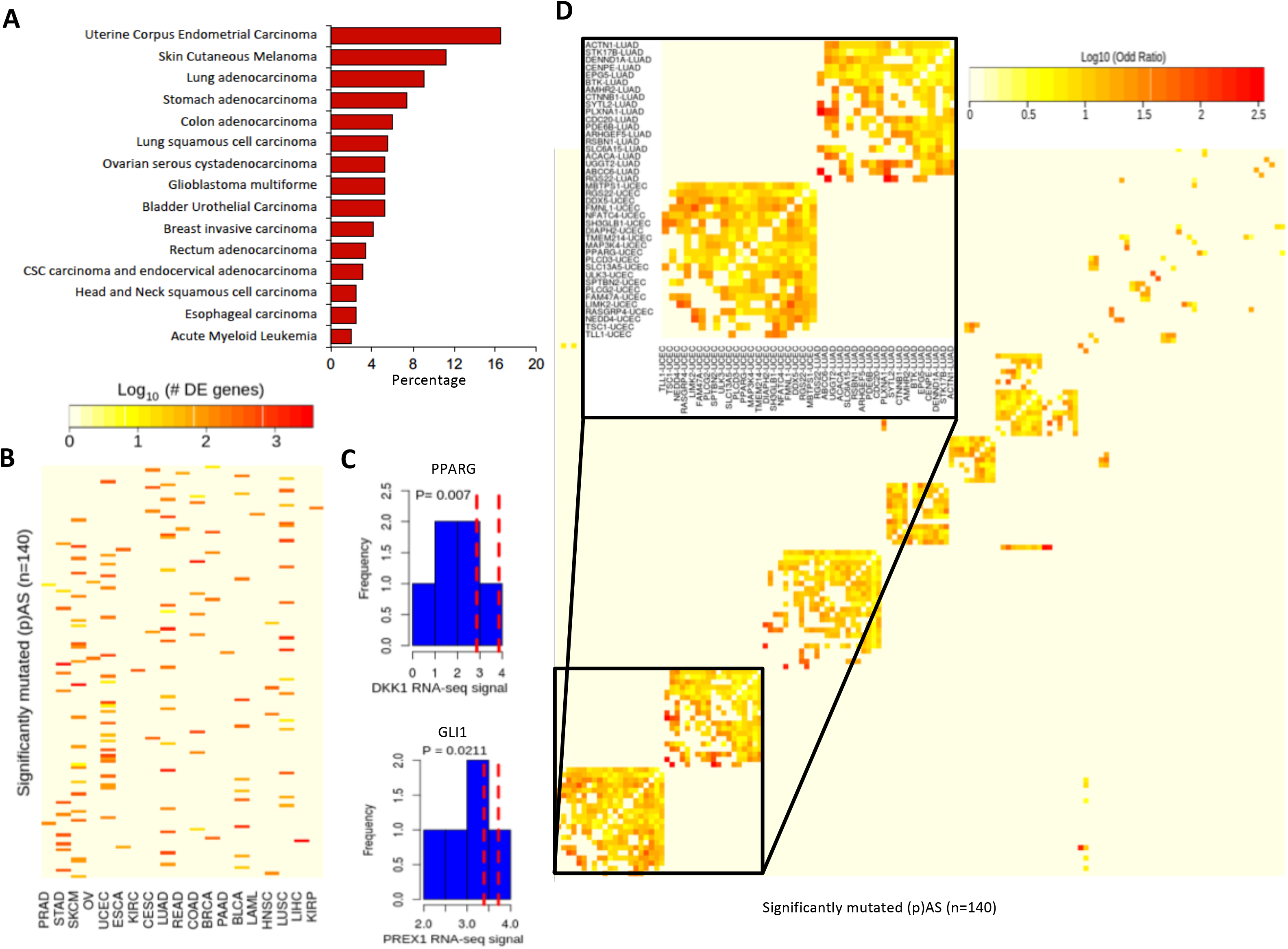
Recurrently mutated (p)AS are associated with distinct molecular signatures (A) Frequency distribution of recurrently mutated (p)AS among different cancer types. The 15 cancer types with the highest number of recurrently mutated (p)AS are shown. (B) Matched RNAseq data from 19 cancer types revealed that mutations in (p)AS of 140 different proteins are associated with more than 10 differentially expressed genes (FDR<0.05). The number of differentially expressed genes is color coded in log scale. (C) RNAseq signals for DKK1 in endometrial carcinoma and PREX1 in stomach adenocarcinoma are shown. DKK1 and PREX1 were identified as top differentially expressed genes with highest log fold changes when comparing RNAseq data for mutations within and outside the pAS of PPARG and GLI1, respectively. The red lines represent signals for samples with mutations in the pAS of PPARG and GLI1. (D) Overlap between differentially expressed genes as a result of mutations in the (p)AS of 140 proteins. Overlapping differentially expressed genes were identified using Fisher’s exact test and clustered based on log odd ratios of the overlap. The heat map reveals that cancer-specific mutations located in (p)AS lead to overlapping gene expression changes.

Somatic missense mutations can disrupt inhibitory allosteric interaction and, as a result, lead to increased activity of the mutated protein. For 12 genes with significant mutated (p)AS (PTPN11^55^, PDGFRA^56^, KIT^57^, AKT3^24^, Vav1^58^, BLK^59^, CARD11^60,61^, FLT3^62^, NLRP1^63^, NOTCH2^64^, EGFR ^65^ and SPATA13[ASEF2]^66^ PLCG1^67^), we found literature support for activity increasing effects of somatic cancer missense mutations in (p)AS. For other genes, there exists indirect support for altered allosteric inhibition, as is the case of PANX1(**Fig. 4D**). Pannexin −1, the protein encoded by PANX1, forms a hexameric plasma membrane channel that in the active form releases ATP in the extracellular space. A truncation mutation, which is found enriched in metastatic breast cancer, enhances Pannexin-1 channel activity and drives metastatic efficiency, likely due to the removal of the autoinhibitory switch in the truncation mutant^68,69^. Cancer missense mutations in this autoinhibitory switch of PANX1 may promote metastatic activity in a similar way. Although missense mutations in (p)AS can weaken intramolecular regulatory interactions, they may also strengthen them or affect interactions with allosteric effectors and as a result change protein activity, localization or stability. An apposite example is the well-known oncogenes b-catenin (CTNNB1), which we find to have recurrent cancer missense mutations in its pAS. Consistent with the prediction of the AS, structural and binding studies have revealed that the N-terminal tail (and C-terminal tail) of b-catenin transiently interacts with other parts of the protein and, thereby, modulates interactions with binding partners^70–72^. However, the N-terminal tail also harbors multiple phosphorylation sites that are key to the regulation of b-catenin’s ubiquitin-mediated degradation and subcellular localization, and the oncogenic potential of missense mutations located in this N-terminal tail has been explained mainly by changes in protein stability and localization^73,74^.

To investigate the potential impact of missense mutations in (p)AS more broadly and search for molecular signatures associated with these mutations, we decided to exploit RNA sequencing data (RNAseq) from NIH-GDC^54^. RNAseq data from 19 cancers showed that in 140 instances mutations found in the (p)AS of 126 of the 302 genes are associated with more than 10 differentially expressed genes (FDR<5%; **Fig. 5B**). 49 of these differentially expressed genes sets are significantly enriched in cancer or cancer pathway annotations (**Supplementary Table 4).** Moreover, detailed analyses of the RNAseq data revealed so-far unknown connections between somatic cancer missense mutations and molecular signatures and provide new insights into unknown or less-studied mechanistic relations. For instance, we selected all transcription factors (NFATC4, GLI1, PPARG and LARP7) among the genes that have differentially expressed genes associated with mutations in their (p)AS and investigated whether the differential expression is, at least in part, the result of increased transcriptional activity of these transcription factors. To this end, we assessed co-expression (Pearson correlation) of these transcription factors and their literature-curated target genes. For all four transcription factors, co-expression is significantly higher (T-test, **Supplementary Table 5)** in patients with mutations inside (p)AS when compared to those with mutations outside the p(AS) of these transcription factors. In the specific case of the PPARG gene, which encodes the peroxisome proliferator-activated receptor gamma and plays an oncogenic role in different cancers^75–77^, missense mutations in its pAS in endometrial carcinoma are associated with significantly increased RNA expressions of DKK1 (T-Test, P < 7.0E^−3^, **Fig. 5C**). Dickkopf-1 (DKK1) is a secreted Wnt/β-catenin pathway antagonist, and high expression levels of DKK1 are associated with shorter survival in various cancers, likely due to its effect on cancers’ metastatic behavior^78,79^. Intriguingly, elevated levels of PPARG have recently also been associate with more aggressive metastatic prostate cancer behavior and poor prognosis^77^. The mechanistic relation between mutations in the pAS of PPARG and the increased expression of DKK1 uncovered here may link the effect of both genes on the metastasizing behavior of certain cancers. In the case of the transcription factor GLI1, missense mutations in its pAS in stomach adenocarcinoma are associated with significantly elevated RNA levels of PREX1 (T-Test, P < 2.1E^−2^, **Fig. 5C**). GLI1 and PREX1 have been associated with mechanisms that sustain cancer stem cells^80,81^. Autocrine VEGF/neuropilin-2 signaling is critical for sustaining cancer stem cells in different cancers, and neuropilin-2 signaling has recently been shown to lead to the PREX1-mediated expression of ERK, which underlies the formation and function of cancer stem cells. Importantly, VEGF/neuropilin-2 signaling also contributes to the induction of GLI1 expression, which enhances neuropilin-2 expression and thus creates the autocrine loop. The link between GLI1 and PREX1 expression, due to mutations in GLI1’s pAS, provides new insights into how the autocrine loop in the VEGF/ neuropilin-2 signaling sustains stem cell properties in cancers.

An analysis of the similarity between differentially expressed genes sets associated with mutations in (p)AS of gene pairs revealed a pronounced concordance in expression changes within certain tumor types (**Fig. 5D**). An enrichment analysis revealed that the sets of overlapping differentially expressed genes in uterine corpus endometrial carcinoma, melanoma, stomach adenocarcinoma, bladder urothelial carcinoma, and lung squamous cell carcinoma are significantly associated with cancer pathways (**Supplementary Table 6)**. The concordance in expression changes as a result of mutations in (p)AS of different genes suggests the existence of potential functional relationships. Indeed, for some of the gene pairs that display concordance in expression changes, mechanistic relationships are well known such as the regulation of PPARG expression by NFATC4^82^, the regulatory impact of CDC20 on CTNNB1^83^ or of CTNNB1 on CCND2^84^. Moreover, some differentially expressed genes do not only overlap between pairs of genes with mutated (p)AS but multiple such genes. Nearly half (45%) of the genes with bladder urothelial carcinoma-associated mutations in their (p)AS change the expression of the genes SAA1, NT5E and KRT14. KRT14 has recently been shown to mark a subpopulation of bladder basal cells that play a pivotal role in tumorigenesis^85^ Thus, our analysis reveals that mutations in (p)AS are associate with specific molecular signatures of cancers.

### Molecular simulations validate altering effects of missense mutations in AS

Missense mutations in AS can enhance tumorigenesis via multiple effects including the direct weakening (in oncogenes) or strengthening (in tumor suppressors) of autoinhibition and the modulation of the interaction with allosteric effectors. We investigated the molecular effects of missense mutations in three genes with significantly mutated AS for which structures of switches are available: ASK1, DAPK2 and EIF4G1. ASK1^86^ and DAPK2^87^ have been shown to have tumor suppressor effects and missense mutations are expected to reduce activity, while EIF4G1 promotes the translation of malignancy-associated mRNAs^47^ and, thus, missense mutations in its AS are expected to enhance activity. Microsecond molecular dynamics (MD) simulations of wild-type and mutant sequences revealed molecular changes that are consistent with these hypotheses.

ASK1 (MAP3K5) belongs to the apoptosis signal-regulating kinases that are part of the p38 and JNK MAP kinase pathways. Inactivating mutations in ASK1 contribute to the development of melanoma^86^. A recent study determined the structure of the AS in Ask1, which consists of seven tetratricopeptide repeats and a pleckstrin homology domain (**Fig. 6A**), and revealed that the mutation R395E opens the compact structure of this AS, thereby decreasing kinase activity^88^. Large-scale MD simulations of AS mutations R300W and V591M, which were found in esophageal carcinoma, demonstrate opening of the compact structure of the autoinhibitory switch, as monitored by distances between residues in its tetratricopeptide repeat core (**Fig. 6A, right**), which suggests that they may result in a loss of kinase activity similar to the R395E substitution studied previously.

**Figure 6.**
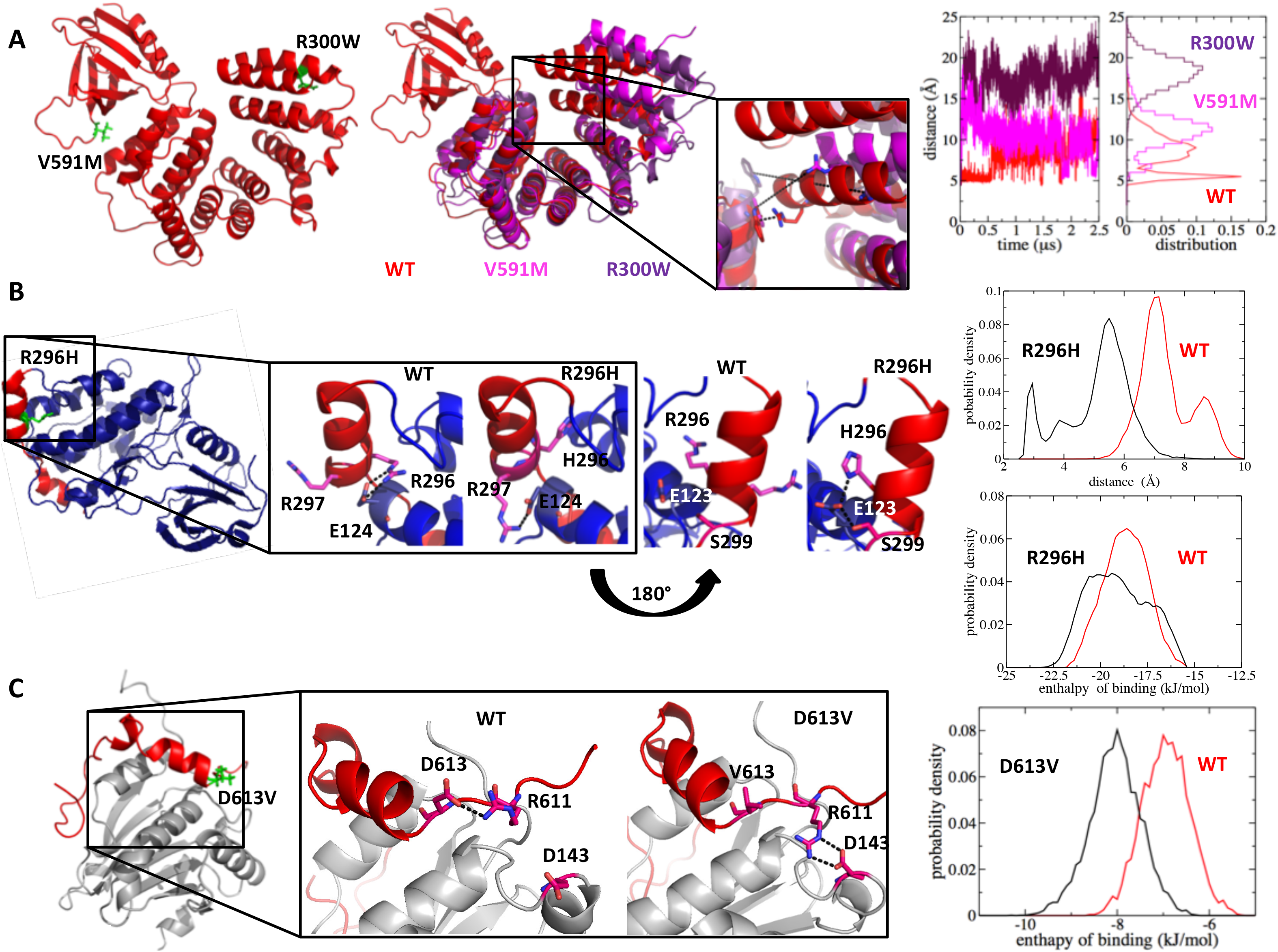
**Molecular dynamics simulations of wild type and mutated allosteric switches.** (A) Left, allosteric switch of wild type ASK1 in ribbon representation (red) with mutated residues R300 and V591 highlighted in green. Middle, representative structures of the wild type (red) and mutated (magenta and purple) allosteric switches after 2.5 µs MD simulations. Structures were aligned on the PH domain (top left), to help visualizing the opening of the tetratricopeptide repeat core in the mutants. Right, distances between residues R322 in W509 and the corresponding distributions, which were used to monitor compactness of the allosteric switch. (B) Left, ribbon representation of wild type DAPK2 with kinase domain in blue and allosteric switch in red. The mutated residue R296 is highlighted in green. Middle, structural detail of the interface between kinase domain and allosteric switch after 2.5 µs of MD simulations of wild type and R296H mutant. Right, distribution of the distance between E123 and R296 (H296) in wild type and mutant simulation. Distribution of enthalpy of binding between kinase and switch for wild type and mutant simulation. (C) Left, ribbon representation of the allosteric switch of wild type eIF4G1(red) in complex with eIF4E (grey). The mutated residue D613 is highlighted in green. Middle, structural detail of the interface between eIF4G1 and eIF4E after 2.5 µs of MD simulations of wild type and D613V mutant. Right, distribution of enthalpy of binding between eIF4G1 and eIF4E for wild type and mutant simulation.

DAPK2 is a pro-apoptotic Ca^2+^/calmodulin (CaM) regulated serine/threonine kinase that, like its closest homolog DAPK1, is a tumor suppressor^87^. Phosphorylation of S299 in its autoinhibitory switch places negative charges next to E123 and, thus, leads, most likely as a result of disrupted autoinhibition, to increased kinase activity^89^. MD simulations reveal that the uterine corpus endometrial carcinoma-associated R296H mutation changes the salt-bridge network and enables new interactions between the H296 and E123 (**Fig. 6B**), which may reduce the activating effect of the phosphorylation of nearby S299. Moreover, energy calculations suggest that the mutation may strengthen interactions between the inhibitory AS and the kinase domain (**Fig. 6B, right**). Together, these analysis results of the wild type and R296H mutation simulations are consistent with the hypothesis that the R296H mutation reduces activity/activation of DAPK2.

eIF4G1 is, as mentioned before, a eukaryotic translation initiation factor with helicase activity that is inhibited by an AS^47,49^. Binding of the oncogenic protein eIF4E to the AS of eIF4G1 relieves autoinhibition and promotes mRNA translation. The glioblastoma-associated D613V mutation is located at the interface between eIF4G1 and it allosteric effector eIF4E. Microsecond simulations of the eIF4G1- eIF4E complex suggest that this mutation leads to the loss of intramolecular interactions between eIF4G1 residues R611 and V613 as well as the gain of new intermolecular interactions between R611 in eIF4G1 and D143 in eIF4E (**Fig. 6c**). Moreover, the D613V mutation is likely to stabilize the interaction between eIF4G1 and eIF4E as suggested by calculations of the enthalpy of binding (**Fig. 6C, right**). This stabilization of the interaction is consistent with the hypothesis that the D613V mutations enhances the relief of autoinhibition and, thus, increases helicase activity.

## DISCUSSION

Autoinhibition is a prevalent allosteric regulatory mechanism in signaling proteins because it prevents spurious pathway activation and enables signal transmission only upon integration of appropriate inputs^20–22^. Therefore, it does not come as a surprise that pronounced effects on cell signaling and regulation, including effects that contribute to cancerous cell transformation and progression, have been observed when autoinhibition is modulated via mutations in individual signaling proteins^19,57,58,90–92^. Our systems-level analysis now reveals that cancer fusion breakpoints and cancer-associated missense mutations are found with significant enrichment within inhibitory allosteric switches across cancer genomes. Importantly, the enrichment is for fusion breakpoints tied to oncogenic activity, consistent with the expected effect of removal or truncation of an inhibitory allosteric switch. This “specificity” in the enrichment of cancer fusion breakpoints within switches of oncogenes, thus, not found in tumor suppressor genes or fusion proteins associated with non-cancer diseases, highlights the potential predictive benefits of integrative studies that combine protein regulation information with cancer genomics data. Our finding that missense mutations rates are higher within allosteric switches compared to outside may be perceived as non-surprising in the light of the previous observation that cancer mutations are enriched at or close to phosphosites^30,31^ and the fact that inhibitory allosteric switches often contain phosphosite clusters that are involved in the regulation of switch function^22,28^. However, cancer mutation rates at or close to phosphosites are significantly higher within allosteric switches compared to outside, which demonstrates that not all phosphosites are equal and that particularly those within allosteric switches are altered in tumor cells. Allosteric switches often act as signal integrating platforms where multiple allosteric effectors contribute to signal transmission, also via (de)phosphorylation or mediated via phosphorylation^93^. Thus, the multi-functionality of these integrative platforms may make them more susceptible to interference that provides selective advantages to cancer cells.

We exploited the enrichment of cancer missense mutations and fusion breakpoints in allosteric switches for the identification of potential cancer driver genes, which follows the rational of previously developed methods that assess the compositional bias of somatic cancer mutations in human genes^30,32–37^. However, our approach complements existing ones as it focusses specifically on protein regulatory sites, which are often short and located in intrinsically disordered regions and as such missed by other approaches. As a result, we are also able to identify potential drivers that have only few somatic gene alterations in regions hat encode short allosteric switches (**Supplementary Fig. 8**). When using cancer fusion data, we identify predominantly known cancer drivers and genes that have been associated with cancer development or progression previously, lending confidence to the overall approach and the driver potential of the selected genes that have not been studied in the cancer context before. Moreover, identified known drivers are nearly all, as expected, oncogenes. It is important to note that different mechanisms can make a fusion gene a driver, including changes in expression, localization or activity, the latter most often due to oligomerization and the removal of inhibitory allosteric switches^6,11,53^. It has been demonstrated that removal of inhibitory switches can be sufficient to confer cell transforming potential to certain drivers but that such removal is not absolutely necessary^39^. Autoinhibition can even be overcome by enforced oligomerization^39^. Thus, our approach will miss drivers that have inhibitory switches but get activated because they oligomerize as a result of gene fusion and not because the switch is removed or shortened. For those cases where the fusion product has dimerization domains as well as an altered allosteric switch, oligomerization and reduced autoinhibition could synergistically enhance oncogene activity as demonstrated for individual cases previously^53^.

Genes with significant enrichment for somatic cancer missense mutations in allosteric switches are also strongly associated with cancer development and contain numerous established drivers. Predicting the molecular phenotype that results from these mutations is, however, more complex than predicting the one that results from switch disruption in gene fusions. Mutations in allosteric switches can weaken or strengthen intramolecular regulatory interactions, thereby increasing or reducing protein activity, or modulate interactions with allosteric regulators^61,91^. Indeed, biophysical simulations suggest that cancer-associated missense mutations in the AS of the tumor suppressors ASK1 and DAPK2 reduce activity of these genes, while they increase the activating interactions between that the oncogenes eIF4E and eIF4G1. It is important to note that missense mutations, as well as truncations as a result of gene fusion, in signal-integrating ASs can also affect other aspects of protein regulation than allostery such as cell localization or protein half-life^74,94^, which in turn can promote cancer development. Besides providing leads for the potential molecular effects in oncogenes or tumor suppressors, our analysis demonstrates that missense mutations in allosteric switches are associated with specific transcriptional signatures of cancer. These include cancer-specific and concordant gene expression changes such the concomitant expression modulation of KRT14 as a result of bladder urothelial carcinoma-associated mutations in allosteric switches of multiple genes and specific regulatory relationships between mutated transcription factors and cancer genes.

The presented approach for the identification of potential drivers has some limitations. First, as it relies on annotations for allosteric protein regulation and as these are sparse, we also exploited predictions of allosteric switches. Thus, predictions of drivers can only be as accurate as the ones of allosteric switches. However, new proteomics methods for the mapping of structural changes that occur in signaling cascades have been developed recently that may provide new high-throughput means to identify allosteric switches and help improve predictions of these^95^. Second, we only used cancer fusions and missense mutations, excluding all other types of mutations, which precludes a detailed comparison with other driver-identification approaches that include all types of mutations. Other mutation types can easily be included in the future. Finally, cancer tissue specific gene expression and splicing will determine the functional relevance of a putative driver and the presence of the allosteric switch in it. Thus, predictions need to be combined with cancer specific transcriptomics data to assess biological relevance further.

In summary, our study reveals that cancer fusions and missense mutations have a statistical preference for proteins with inhibitory allosteric switches, respectively the residues therein, which demonstrates that altered autoinhibtion is positively selected for by cancer cells. We show that selection of genes with significantly mutated allosteric switches provides a fruitful way to identify cancer drivers. This new way of identifying drivers is particularly attractive in the light of recent successes in the design and clinical use of precision medicines that stabilize autoinhibited states of oncogenes^25,26,96,97^.

## METHODS

### Data collections and processing

Proteins with experimentally-characterized inhibitory allosteric switches (AS) were identified in a literature search as described previously^22,28^. Boundaries of AS were taken from the original papers in which either assays on deletion constructs or detailed structural characterizations were used to map autoinhibitory protein regions. 234 human proteins with AS were found in the literature search. Proteins with predicted AS (pAS) were identified by using the predictor Cis-regPred^28^. Cis-regPred identifies intrinsically disordered allosteric switches based on protein sequence information as well as publically available proteomics data. Features that Cis-regPred uses to identify pAS include, among others, the amino acid compositional bias, intrinsic disorder prediction scores, enrichment for splicing sites and post-translational modifications of autoinhibitory regions. In addition, electrostatic complementarity between the allosteric switch and other parts of the same proteins is used. Cis-regPred has been trained and validated on independent sets of proteins with autoinhibitory regions that have been mapped by experimental means. Although trained exclusively on inhibitory allosteric switches (autoinhibitory protein regions), Cis-regPred has been shown to identify also activating allosteric switches. This said, most of the pAS identified by Cis-regPred are likely to be inhibitory in nature. Intrinsic disorder predictions scores from Cis-regPred were also used to identify intrinsically disordered protein regions.

Fusion genes were collected from the databases FusionCancer^98^ and ChimerDB 3.0^99^ as well as from Yoshihara *et al*^100^. Only fusion genes were considered in which the fusion breakpoint is within the protein coding region. For this analysis, genomic co-ordinates of fusion proteins breakpoints were mapped to protein co-ordinates from annotation file downloaded from Ensembl^101^. In addition, to identify functional fusion proteins, only fusion proteins were considered with functional domain(s). Domains were identified based on annotations provided by PFAM^102^ and InterPro^103^. Overall, 12’957 fusion genes were collected, and 5’503 in-frame fusion proteins identified that were found in cancer cells and had the fusion breakpoint within protein coding regions. 756 proteins were identified that occurred only in non-cancer fusions.

7’799 cancer mutations and 17’549 phosphorylated residues for the initial enrichment analysis were downloaded from COSMIC database^29^ and phosphositePlus^104^. Gene expression data (level 3), 18’845 somatic mutations for the identification of recurrently mutated (p)AS, and mapped read count data were downloaded from NCI-Genome Data Commons^54^.

### Fusion breakpoint and cancer mutation enrichment analysis

The hypergeometric test was used to assess the enrichment of breakpoints/junctions of fusion genes within (p)AS. Thus, the number of breakpoints disrupting (p)AS regions was compared to the number of breakpoints outside the region, using the number of residues in and outside of (p)AS for normalization. In a second enrichment analysis, the boundary of the (p)AS was extended by including the linker between the (p)AS and the regulated functional domain of each protein. This was done as breakpoints may not fall within the (p)AS region but between the (p)AS and the regulated domain, thus resulting in the removal of the (p)AS but keeping the functional domain intact. For known AS, boundaries of AS and the regulated domain are clearly defined, which enables identification of the “extended AS”. For predicted AS, it is not a priori clear which domain or sequence region is regulated by the (p)AS, particularly for multidomain proteins. Therefore, the following rules were applied. 1. For single domain proteins with pAS, the unique domain present was considered the domain that is regulated by the pAS. 2. For multidomain proteins, only those proteins were considered that have a kinase, DNA-binding or guanine nucleotide exchange factor domain. These domains were assumed to be regulated by the pAS.

To assess the enrichment of for COSMIC cancer mutations within (p)AS, the hypergeometric test was applied as explained for the fusion breakpoints. Averaged mutations rates per residue inside and outside of (p)AS were compare using the T-test.

### Identification of genes with significantly altered (p)AS

For the identification of significantly mutated (p)AS, 18’845 somatic mutations from 10,177 patients were downloaded from NCI GDC^54^. Cancer-specific missense mutations were mapped to (p)AS regions using BEDtools^105^. Then P-values of the binomial probabilities of finding k or more missense mutations within regions that encode the (p)AS were calculated under the assumption that all residues have the same probability of being mutated. Thus, given the total number of mutations in a protein, its length and the length of the (p)AS, one can calculate the binomial probabilities of finding k or more missense mutations within regions that encode the (p)AS under the null hypothesis that the probability of mutations is equal across the gene, an approach also used by e-Driver^36^. Consistent with the null hypothesis, no significant difference in the rate of coding-silent mutations inside and outside of (p)AS was detected (**Supplementary Fig. 9**). Moreover, several algorithms have recently been used and compared in the identification of cancer drivers, including e-Driver^106^. Different quality metrics were applied to compare methods and eliminate outlier predictions. One of these metrics is the divergence of observed from theoretically expected P vales, which, when large, may indicate that a methods statistical assumption is not well satisfied. E-Driver produced less often outliers, when analyzing cancer-specific data, than many of the other established driver-identification methods, thus providing confidence for the statistical approach used. Once P values for all (p)AS in each gene of a specific cancer were calculated, the Benjamini–Hochberg false discovery rate algorithm was applied to correct for multiple testing.

For fusion breakpoints, the same procedure was used but missense mutations replaced by breakpoint locations and (p)AS replaced by extended (p)AS (for extended (p)AS definition see above). A cancer-specific analysis of fusion data did not provide any hits for significantly altered (pAS), thus a pan-cancer analysis was carried out instead. Genes with significantly altered (p)AS as a result of gene fusion were identified and ranked after correction for multiple testing using the Benjamini–Hochberg false discovery rate algorithm.

### Dimerization domain analysis

A list of dimerization domains was compiled based on PFAM^102^ and InterPro^103^ annotations. These dimerization domains were then mapped to the fusion proteins and assed how often fusion involving proteins with (p)AS lead to the aaddition of a new dimerization domain.

### Differential expression and downstream analysis

Mapped raw reads count data downloaded from NCI-Genome Data Commons ^54^ was used for the identification of genes differentially expressed between patients with mutations inside and outside the (p)AS. Patients having mutations within as well as outside the (p)AS region were removed from the analyses. Further, only those mutations were selected for which at least 2 replicates exist. This was done to reduce the variance in the data and improve the identification of differentially expressed genes. This selection criteria resulted in 79 BLCA, 33 BRCA, 31 CESC, 83 COAD, 12 ESCA, 33 HNSC, 4 KIRC, 4 KIRP, 5 LAML, 90 LIHC, 165 LUAD, 137 LUSC, 10 OV, 5 PAAD, 16 PRAD, 8 READ, 297 SKCM, 78 STAD and, 306 UCEC tissues/samples which have matched count and mutation data. After initial filtering, DESeq2 was used to identify differentially expressed genes by taking adjusted P-value of << 0.05. Further, we only considered genes with recurrently mutated (p)AS that have at least 10 differentially expressed genes. These filtering result in 140 gene-cancer combinations consisting of 126 unique genes in 19 cancer tissues. Enrichment of differentially expressed genes was done by EnrichR^107^ and PathwAX^108^. Then the overlap of these sets of differentially expressed genes within cancers was determined. This was done by iterating the differentially expressed (DE) genes starting from top 10% of genes in each cancer followed by 10% increment in each iteration till all the genes were covered (100%). Each iteration in a cancer set was compared with iteration in other cancer set thus, resulting in 10 × 10 steps. For each steps log odd ratio and p-value was calculated using Fisher’s Exact test. For clustering log odd ratio of most significant (lowest) p-value was extracted for each pair-wise overlapping DE genes set. their log odd ratio calculated and Fisher’s exact test applied to find the significance of overlap.

### Expression correlation between transcription factors with significantly mutated (p)AS and their target genes

(p)AS containing transcription factors (TFs) that are enriched in somatic cancer missense mutation were identified as described above. Mutations mapping to these TFs were mapped to corresponding gene expression data sets (FPKM values) that were downloaded from NCI GDC. For the mapping, unique IDs of mutations and RNA-seq expression data were matched. The effect of mutation on TF activity was assessed by comparing expression data from patients with mutations within and outside the (p)AS. Targets of transcription factors involved in this study was downloaded from GTRD database^109^. Once all the targets of TFs were downloaded gene expression profile based co-expression between TF and target genes was calculated as Pearson Correlation Coefficient (PCC). P-values of co-expression (PCC) were calculated using “T-test”. To find the significant change in PCC between mutations within AI/CRE region to mutation outside AI/CRE region, T-test was applied.

### Statistical analyses

Statistical analyses were performed using the JAVA apache maths library “R”. Gene Ontology and pathway enrichment analyses were done using EnrichR^107^ and PathwAX^108^.

### Molecular dynamics simulations

We performed 2.5 µs molecular dynamics (MD) simulations for each studied system of both wild-type and mutants. Prior MD production, missing electron loop densities of crystal structures were reconstructed using Rosetta loop remodelling. Next, protonation states for all titratable residues in the resulting structures were determined. The effective pKa for each titratable group was computed using the WHATIF pKa calculation package as described previously^110^. The linearized Poisson-Boltzmann equation was computed using a dielectric constant of four for the protein using Adaptive Poisson-Boltzmann electrostatic solver^111^. Molecular dynamics simulations were carried out using the modified AMBER ff14SB force field^112^. The simulations began with 5000 steps of steepest descent energy minimization were followed by the reassignments of velocities from the Maxwell distribution at 300 K every 1 ps for 1000 ps, and by a final equilibration of the system for 2.0 ns. After equilibration, the molecular dynamics trajectories were integrated using the GPU-accelerated *pmemd* engine in Amber14. The simulations were conducted using the isobaric-isothermal ensemble^113^ at 300 K and 1 atmosphere and using long-range non-bonded interactions with a 12 Å residue-based cutoff. Long-range electrostatic forces were calculated using the particle-mesh Ewald sum^114^. Bonds to hydrogen atoms were maintained with the SHAKE algorithm^115^ and an integration step size of two femtoseconds was used.

## ACKNOWLEDGEMENTS

We thank the Canadian Institute of Health Research (CIHR), Genome Canada, Genome BC and the Michael Smith Foundation for Health Research (MSFHR) for their valuable support. This research was supported by Basic Science Research Program through the National Research Foundation of Korea (NRF) funded by the Ministry of Science, ICT & Future Planning (NRF-2014R1A1A1003444).

